# Sampling in structure-token space enables accurate prediction of multiple protein conformations

**DOI:** 10.64898/2026.03.03.708411

**Authors:** Zhiyuan Wang, Yue Yu, Wei-Mou Zheng, Chungong Yu, Dongbo Bu

## Abstract

Proteins often function through multiple conformations, yet most structure-prediction methods are biased toward a single dominant state. Here we present MSFold, a training-free framework that explores the discrete, sequence-conditioned structure-token space of ESM3 using parallel tempering, coupling high-temperature exploration with low-temperature refinement to cross barriers between conformational states. Across 312 multi-conformation protein pairs, MSFold recovered both conformations in 51.6% of cases, compared with 38.1% for MSA Cluster and 37.8% for AlphaFold3. This performance advantage persisted on a post-ESM3-cutoff evaluation set, arguing against simple structural memorization. We further introduce sequence log-likelihood, a reference-free confidence score based on sequence–structure consistency, which improves ranking of sampled conformations. Together, these results provide a practical means of accessing the conformational information encoded by protein language models and establish MSFold as an effective framework for multi-conformation prediction.

## 1 Introduction

Proteins often execute their biological functions through multiple structural conformations rather than a single static structure, and distinct conformational states underlie ligand recognition, catalysis, transport, and allosteric regulation [1]. Experimental technologies for structure determination can yield essential snapshots of these states; however, the yielded snapshots usually carry incomplete information about conformations: X-ray crystallography and cryo-electron microscopy can favor trapped, stabilized, or averaged structural states [2, 3], and the conformation reconstructions often collapse ensemble information into a single structural state. In addition, nuclear magnetic resonance technology offers advantages in resolving conformational diversity but shows limitations in processing large proteins [4, 5]. As a result, the current structural repository, Protein Data Bank (PDB) [6], provides only a partial view of protein conformational equilibria [7, 8], and alternative conformations remain substantially underrepresented. Hence, predicting multiple functionally relevant conformations from a single sequence is an important goal for structure prediction methods.

Accurate prediction of multiple conformations requires effective exploration of the structure-token space and reliable selection from candidate conformations. In principle, the diverse conformations and their dynamics can be characterized by molecular dynamics (MD) simulations, which, however, are limited by sampling cost, accessible timescales, and force-field accuracy [9]. The deep learning-based prediction approaches, such as AlphaFold [10, 11] and RoseTTAFold [12], were primarily optimized to approximate a single highly probable conformation and often exhibit biases toward the dominant state. Recent advances based on multiple sequence alignment sampling [13], clustering [14], diffusion models [15, 16], and flow matching [17] can increase structural diversity in some cases, but still remain far from achieving accurate prediction across diverse proteins. A further challenge is ranking—the existing confidence scores, such as pLDDT and pTM [10], cannot reliably identify high-quality alternative conformations from the sampled ensembles [18, 19].

Protein language models trained on large and diverse sequence corpora can learn broad representations that transfer to structural and functional tasks [20]. ESM3 [21] extends this paradigm by jointly modeling sequence, structure, and function within a generative token-based framework. Its structure tokenizer encodes a protein structure as a sequence of residue-level discrete structure tokens, each representing the local structural environment around the corresponding residue, while a decoder reconstructs the 3D structure from the token sequence. This discrete representation provides a natural space for exploring alternative structural realizations through structure-token sampling. However, under the iterative sampling schemes considered here, argmax updates and temperature-controlled stochastic sampling tend to remain close to the current token configuration and therefore often explore only a limited region of structure-token space, making it difficult to reach structurally distinct alternative states.

The existing studies have established most of the necessary components for multi-conformation prediction, but an effective sampling strategy has remained elusive. Here, we provide this missing component by introducing MSFold, a training-free framework that applies parallel tempering [22] to sample the structure-token space, together with a confidence score for ranking the resulting conformations. MSFold explores the sequence-conditioned structure-token space using replicas distributed across a temperature ladder: high-temperature replicas promote broad exploration and facilitate barrier crossing, whereas low-temperature replicas preserve structural accuracy. The exchanges between neighboring replicas couple global exploration with local refinement, enabling MSFold to overcome the local-search limitations of standard decoding. Across a benchmark of proteins with experimentally determined multiple conformations, MSFold achieved a higher multi-conformation success rate than existing approaches, including AlphaFold3, primarily through improved recovery of Fold 2 while maintaining high Fold 1 accuracy. Together, these results establish MSFold as an effective tool for multi-conformation prediction.

## 2 Results

### 2.1 MSFold recovers multiple protein conformations across diverse proteins

MSFold takes an amino acid sequence as input and explores the sequence-conditioned structure-token space learned by ESM3 using parallel tempering (Fig. 1a, b). Multiple replicas sample structure-token sequences at different temperatures, with replica exchange allowing information to propagate across the temperature ladder. The sampled token sequences are decoded into an ensemble of protein conformations, which are subsequently ranked using a confidence score. In this study, MSFold uses 40 replicas and performs 500 sampling steps, yielding an ensemble of 20,000 conformations.

**Fig. 1:**
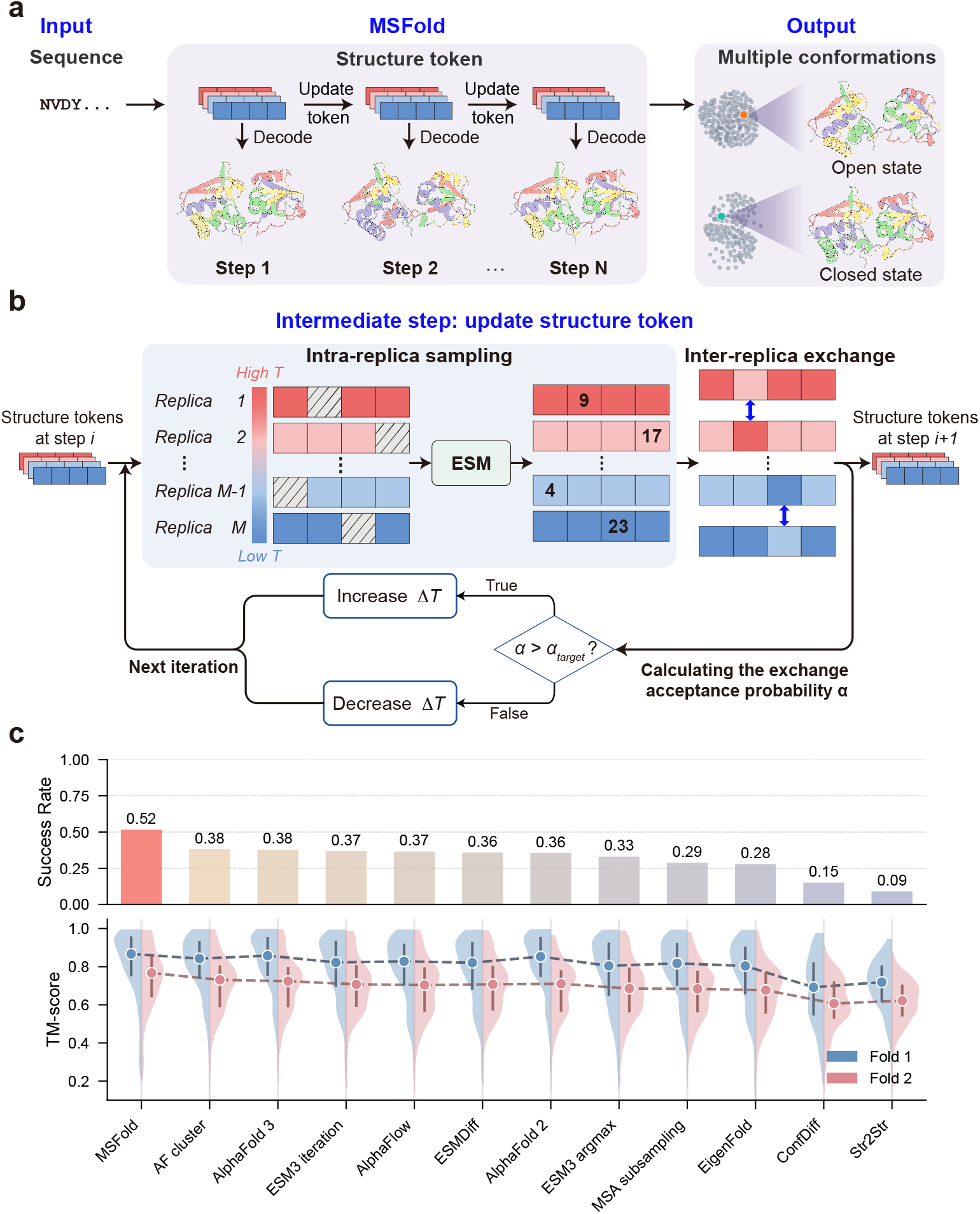
Overview of MSFold and its performance in multi-conformation prediction. **a,** MSFold predicts multiple protein conformations by sampling the ESM3 structure-token space using parallel tempering. It uses 40 replicas of the structure-token sequence distributed along a temperature ladder and performs 500 sampling steps. **b,** At each sampling step, MSFold performs a randomized sweep over all residue positions within each replica, masking and resampling the corresponding local block from temperature-scaled ESM3 structure-token distributions, followed by an attempted exchange of selected tokens between neighboring replicas. The temperature ladder is adaptively adjusted according to the exchange acceptance probability. **c,** Performance on a representative benchmark comprising 312 multi-conformation protein pairs with experimentally determined multiple conformations. MSFold achieves the highest multi-conformation prediction success rate, defined as the fraction of protein pairs for which both Fold 1 and Fold 2 conformations are recovered with a TM-score of at least 0.75.

We evaluated MSFold on a representative benchmark comprising 312 multi-conformation protein pairs with experimentally determined multiple conformations. These protein pairs encompass diverse modes of conformational variation, including apo-holo transitions [23], fold switching [24], and secondary-structure changes (see Supplementary Data for details). We assessed the predicted conformations by quantifying their structural similarity to the experimentally determined conformations using TM-score [25]. Following the convention used in previous work [18], we denote the native conformation preferentially predicted by ESM3 as Fold 1 and the alternative conformation as Fold 2. A target was considered successful if the generated ensemble contained a candidate with TM-score ≥ 0.75 to Fold 1 and a candidate with TM-score ≥ 0.75 to Fold 2.

As shown in Fig. 1c, MSFold achieves the highest multi-conformation prediction success rate among all evaluated methods. Specifically, MSFold correctly recovers both Fold 1 and Fold 2 conformations for 161 out of the 312 protein pairs, resulting in a success rate of 0.516, consistently higher than AlphaFold models (AlphaFold3: 0.378; AlphaFold2: 0.356), generative approaches (AlphaFlow: 0.365; ConfDiff: 0.151; Str2Str: 0.090), and MSA-based approaches (AF Cluster: 0.381; MSA Subsampling: 0.288).

The superior multi-conformation prediction success rate of MSFold mainly stems from its enhanced ability to recover Fold 2 while maintaining high prediction accuracy for Fold 1. Specifically, most evaluated methods show high prediction accuracy for Fold 1: MSFold achieves an average TM-score of 0.821, comparable to AlphaFold3 (0.830) and AlphaFold2 (0.824). The remaining methods achieve Fold 1 TM-scores ranging from 0.679 to 0.800. In contrast, MSFold exhibits a substantial advantage in recovering Fold 2, achieving an average TM-score of 0.740, higher than all other methods; the next-highest scores are ESM3 iterative sampling (0.702), AlphaFold3 (0.701), AF Cluster (0.699), and ESMDiff (0.694) (Supplementary Table S1). This asymmetric performance suggests that existing methods show substantially stronger recovery of Fold 1 than Fold 2.

Overall, MSFold improves multi-conformation prediction primarily by enhancing recovery of Fold 2 while preserving high accuracy for Fold 1, thereby reducing the single-state bias observed in existing methods.

### 2.2 MSFold generalizes beyond known structures

To distinguish generalization beyond the training-data cutoff from prediction on proteins released before the cutoff, we performed two temporal evaluations: one on proteins released after the cutoff to test generalization beyond known structures, and the other on proteins released before the cutoff to examine whether inclusion in the training data improves multi-conformation prediction.

We first assessed whether MSFold generalizes beyond the structures available for ESM3 training rather than relying on memorization of known structures. We identified 33 protein pairs with experimentally determined structures that were released after the temporal cutoff of the ESM3 training data (1 May 2020), forming a temporally held-out evaluation subset. As shown in Table 1, MSFold achieved a success rate of 0.576 on this subset, comparable to its performance on the original test set (0.516). In contrast, most evaluated methods showed reduced performance (ESMDiff: 0.273; ESM3 with argmax decoding: 0.333; ESM3 with iterative structure-token sampling: 0.303). Analysis of average TM-score further confirmed that MSFold maintained its superiority in recovering alternative conformations on this evaluation subset. These results suggest that MSFold generalizes beyond the structures available during ESM3 training and that its advantage is unlikely to be explained by simple memorization of known structures. We next investigated whether having both Fold 1 and Fold 2 reference conformations available before the training cutoff improves multi-conformation prediction. We used AlphaFold 2 [10] as an example and extracted from the original test set 238 protein pairs whose experimentally determined conformations were released before the AlphaFold 2 training cutoff. On this subset, AlphaFold 2 achieved a success rate of 0.370, compared with 0.356 on the original test set and 0.521 for MSFold on the same subset. This result indicates that having both Fold 1 and Fold 2 reference conformations available before the training cutoff alone is insufficient for reliable multi-conformation prediction. Instead, the major limitation likely lies in the prediction objective and inference procedure, which favor a single conformation rather than exploring alternative conformational states.

**Table 1:**
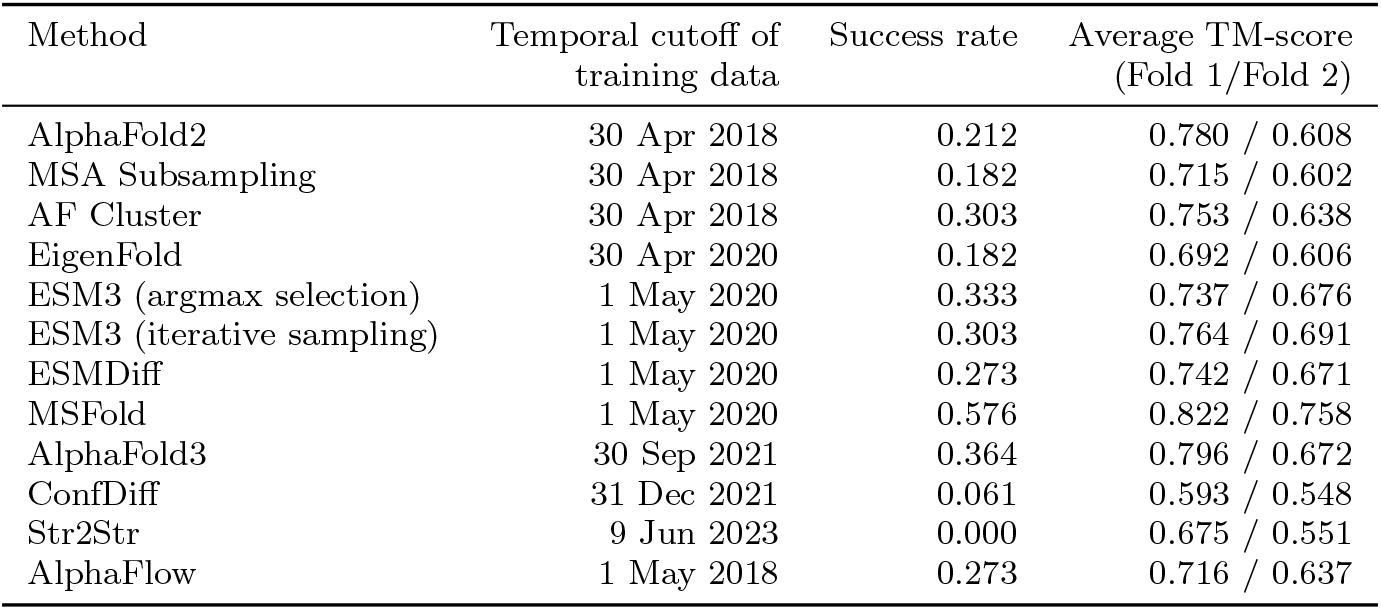
Temporal evaluation on proteins released after the ESM3 training-data cutoff. This table compares the performance of MSFold and existing methods on a temporally held-out evaluation subset comprising 33 protein pairs. The table shows the TM-score of the best-matching conformations in the sampled candidate conformations against Fold 1 and Fold 2 conformations. MSFold achieves the best overall performance, including success rate, and average TM-scores for both Fold 1 and Fold 2.

| Method | Temporal cutoff of training data | Success rate | Average TM-score (Fold 1/Fold 2) |
| --- | --- | --- | --- |
| AlphaFold2 | 30 Apr 2018 | 0.212 | 0.780 / 0.608 |
| MSA Subsampling | 30 Apr 2018 | 0.182 | 0.715 / 0.602 |
| AF Cluster | 30 Apr 2018 | 0.303 | 0.753 / 0.638 |
| EigenFold | 30 Apr 2020 | 0.182 | 0.692 / 0.606 |
| ESM3 (argmax selection) | 1 May 2020 | 0.333 | 0.737 / 0.676 |
| ESM3 (iterative sampling) | 1 May 2020 | 0.303 | 0.764 / 0.691 |
| ESMDiff | 1 May 2020 | 0.273 | 0.742 / 0.671 |
| MSFold | 1 May 2020 | 0.576 | 0.822 / 0.758 |
| AlphaFold3 | 30 Sep 2021 | 0.364 | 0.796 / 0.672 |
| ConfDiff | 31 Dec 2021 | 0.061 | 0.593 / 0.548 |
| Str2Str | 9 Jun 2023 | 0.000 | 0.675 / 0.551 |
| AlphaFlow | 1 May 2018 | 0.273 | 0.716 / 0.637 |

These temporal evaluations demonstrate that MSFold’s performance improvement does not arise solely from the inclusion of known structures in the training data, but from its ability to effectively explore the conformational space encoded by protein language models.

### 2.3 Sampling strategy is critical for multi-conformation prediction

To determine whether the improvement of MSFold arises from enhanced sampling rather than differences in structural representation, we compared MSFold with ESM3 decoded using its standard decoding strategies, including argmax selection and iterative structure-token sampling. In these comparisons, all approaches operate on the same ESM3 structure-token representation and differ only in their sampling strategies.

As shown in Fig. 1c, ESM3 argmax selection and iterative sampling achieve success rates of 0.330 and 0.369, respectively, both substantially lower than that of MSFold (0.516). Detailed analysis reveals that this performance gain is driven primarily by improved recovery of Fold 2 conformations. Although ESM3 argmax and iterative sampling maintain high prediction accuracy for Fold 1, they show substantially reduced accuracy for Fold 2 (average TM-score: 0.679 and 0.702, respectively; Supplementary Table S1). In contrast, MSFold more effectively recovers Fold 2 by enabling exploration beyond the local regions favored by standard decoding strategies. MSFold also outperforms ESMDiff, a fine-tuned ESM3 variant developed specifically for multi-conformation generation using diffusion models [26], indicating that advances in sampling can complement, and in this case exceed, gains obtained through model fine-tuning alone.

These findings identify sampling as a critical determinant of multi-conformation prediction. They reveal that unlocking the conformational information encoded by protein language models requires not only improved representations, but also effective sampling strategies to explore the structure-token space.

### 2.4 MSFold traverses the structure-token space to access alternative states

To understand how MSFold accesses both experimentally determined conformations, we examined the sampling trajectory and resulting conformational ensemble using the LAO-binding protein (LAOBP) as a case study. LAOBP has been experimentally shown to adopt two distinct conformational states, the open state (2LAO) and the closed state (1LAH). For LAOBP, MSFold successfully recovered both states with TM-scores of 0.99 and 0.98, respectively (Fig. 2c). We first characterized the sampled ensemble by projecting the conformations into a two-dimensional space using UMAP [27]. We computed a pairwise similarity matrix of the sampled structures and used 1 – TM-score as the distance metric, with the known native states embedded as reference points. As shown in Fig. 2b, the sampled ensemble forms two well-separated clusters corresponding to the two experimentally determined conformations.

**Fig. 2:**
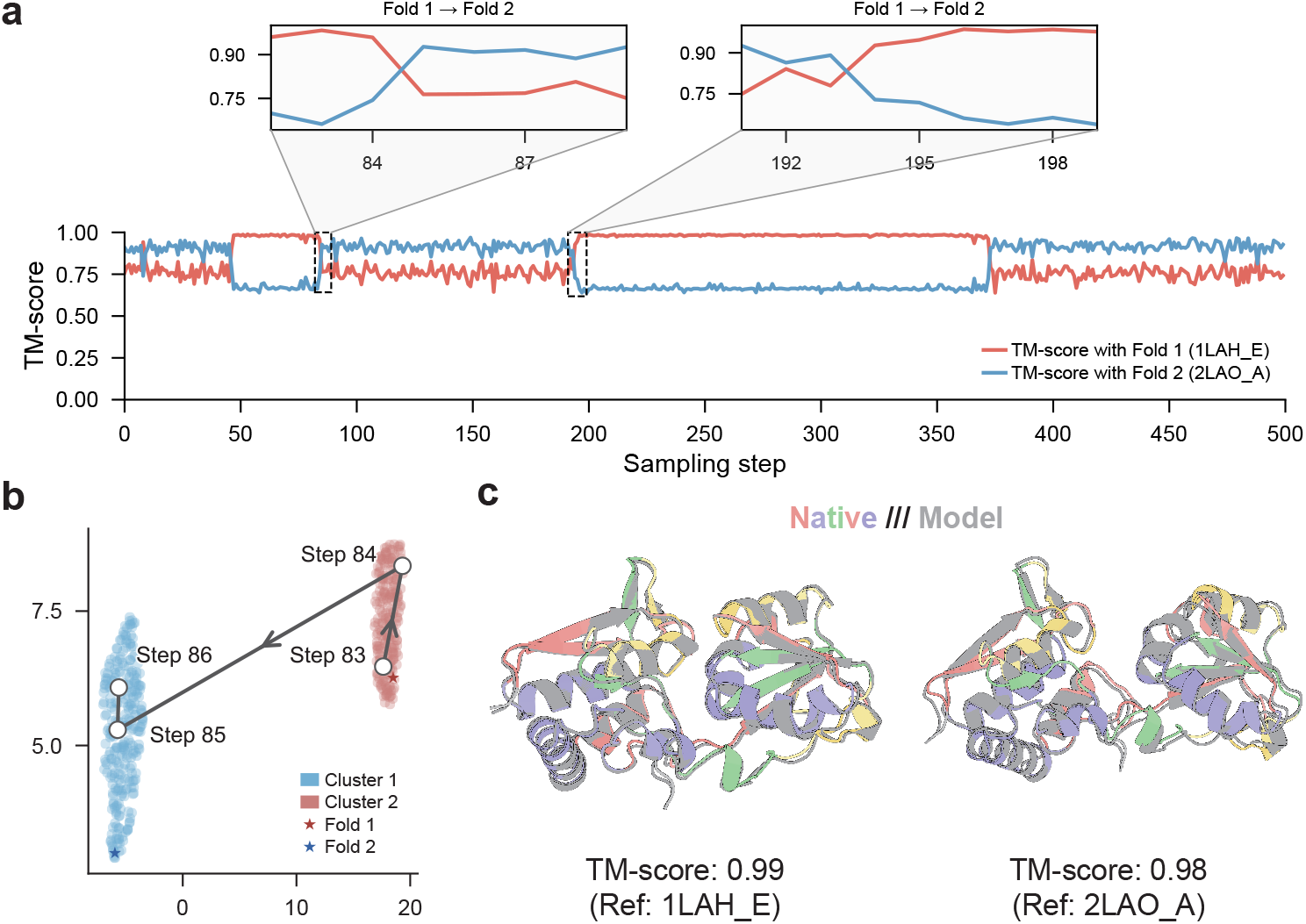
MSFold traverses the structure-token space to access both Fold 1 and Fold 2 states. **a,** Sampling trajectory of MSFold for the LAO-binding protein (LAOBP). We calculated the TM-scores between the sampled conformation and the closed state (Fold 1, 1LAH E; red) and the open state (Fold 2, 2LAO A; blue). The stable lines with only minor fluctuations (e.g., step 1-49) indicate local structural refinements within local regions, whereas line crossings (e.g., step 84-85) indicate transitions between the Fold 1 and Fold 2 state regions. **b,** UMAP projection of the sampled conformation ensemble. The ensemble forms two well-separated clusters that represent two distinct regions corresponding to Fold 1 and Fold 2 conformations. The sampling steps 84 and 85 accomplish a transition between the two regions. **c,** Structural superpositions of predicted models (grey) onto the Fold 1 and Fold 2 structures (colored). MSFold successfully recovered both Fold 1 and Fold 2 conformations with TM-scores of 0.99 and 0.98, respectively.

We next analyzed the sampling trajectory of MSFold for LAOBP (Fig. 2a). At each of the 500 sampling steps, we selected the conformation generated by the lowest-temperature replica as the representative structure and calculated its TM-scores to the Fold 2/open and Fold 1/closed states. These TM-scores are shown as blue and red curves, respectively. A higher red curve than blue curve indicates that the sampled conformation is more similar to the Fold 1/closed state, whereas a higher blue curve indicates greater similarity to the Fold 2/open state. From step 1 to 49, both curves remain relatively stable with only minor fluctuations. During this stage, the sampled conformations consistently resemble the Fold 1/closed state, with TM-scores above 0.80, indicating that the sampled conformations remain close to Fold 1. As sampling proceeds, the two curves occasionally cross, indicating transitions between the Fold 1/closed and Fold 2/open states. For example, the transition from step 84 to step 85 changes from a closed-like to an open-like conformation, demonstrating access to Fold 2/open (Fig. 2a). At step 193, the trajectory returns from the open state to the closed state (Fold 1), enabling further exploration of the conformational space. Two additional case studies, adenylate kinase and catalytic antibody 48G7 Fab, show similar sampling behaviors, as described in the Supplementary Information. Collectively, these findings show two complementary sampling behaviors in MSFold: local refinement around individual conformational states and transitions between distinct conformational states. Local sampling improves structural similarity within a state, whereas state transitions allow the sampler to access other conformational states.

To understand the mechanism underlying transitions between regions, we analyzed how replicas along the temperature ladder explore the conformational space. As shown in Fig. 3a, the low-temperature replica 37 generated a conformational ensemble concentrated in two compact and well-separated clusters, corresponding to localized sampling within individual regions and local refinement. The middle-temperature replica 28 exhibited similar behavior, with differences mainly reflected in cluster sizes. In contrast, the high-temperature replica 11 generated a more diverse ensemble, suggesting broader exploration of the conformational space. The exchange between replicas combines these complementary properties, enabling transitions beyond local regions.

**Fig. 3:**
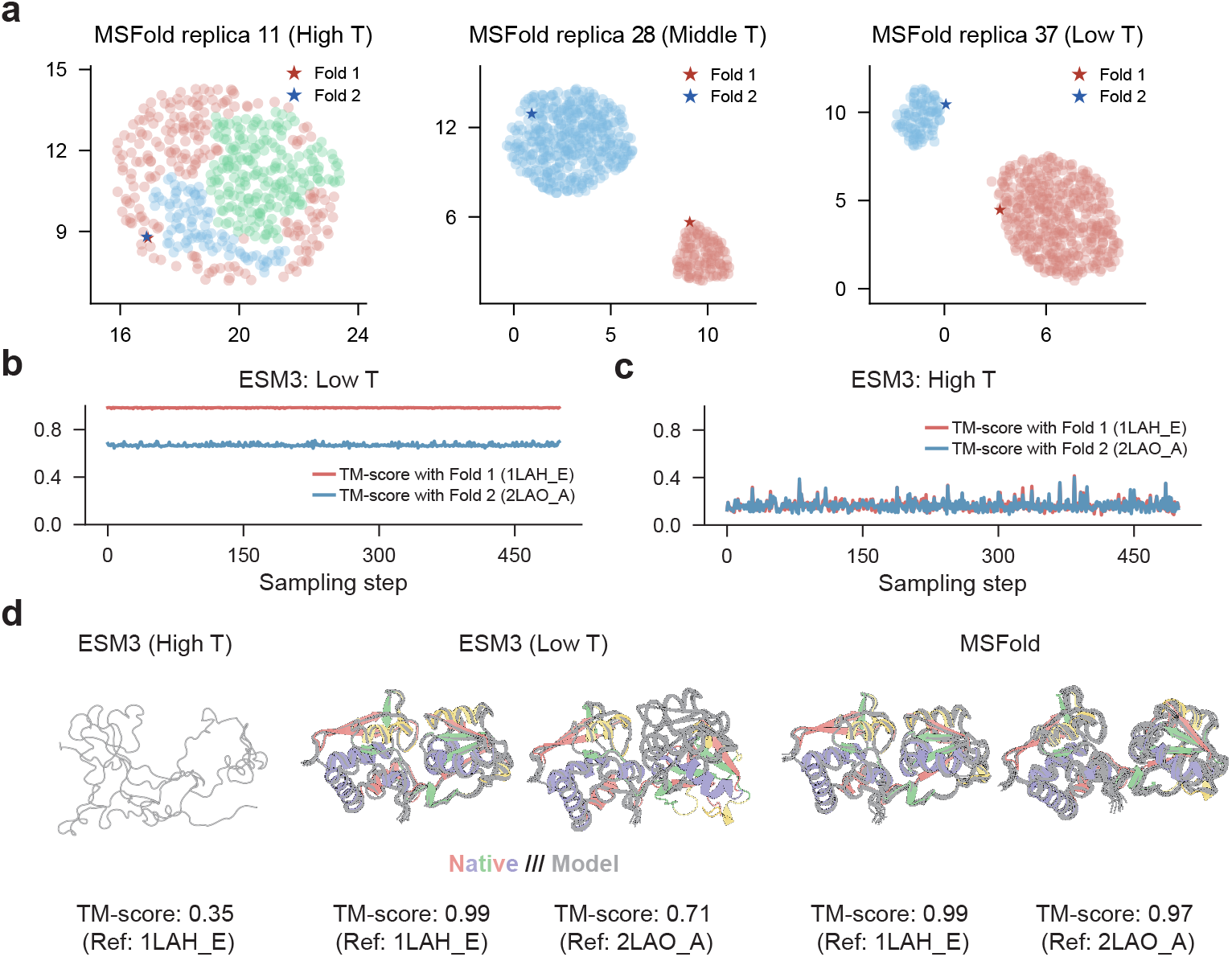
Sampling behavior of MSFold replicas and single-temperature ESM3. **a,** The conformational ensembles generated by MSFold replicas 11 (high *T*), 28 (middle *T*), and 37 (low *T*) for protein LAOBP. Replicas 28 and 37 yield ensembles concentrated in two compact and well-separated clusters corresponding to Fold 1 and Fold 2, whereas replica 11 yields a more diverse ensemble. **b,c,** Sampling trajectories of ESM3 under a single-temperature sampling scheme. At low sampling temperature, ESM3 remains within the Fold 1 region without transitioning to the Fold 2 region. At high sampling temperature, it generates structurally distorted conformations that are similar to neither Fold 1 nor Fold 2. **d,** The best-matching conformations generated by ESM3 and MSFold.

We further performed an ablation study and a single-temperature comparison to validate the role of replica exchange. In the ablation study, we compared MSFold with two variants that either replaced block updates with single-residue updates or disabled replica exchange. Replacing block updates with single-residue updates resulted in only a small change in performance, whereas disabling replica exchange reduced the Fold 1 and Fold 2 TM-scores from 0.840 and 0.792 to 0.802 and 0.775, respectively (Extended Data Table 1), showing that replica exchange contributes to multi-conformation prediction performance. In the second experiment, we compared MSFold with ESM3 under a single-temperature sampling scheme and examined the sampling trajectory of ESM3 (Fig. 3). At high sampling temperatures, ESM3 generated disordered conformations that were similar to neither Fold 1 nor Fold 2 (TM-score ≤ 0.40). In contrast, low-temperature sampling enabled ESM3 to generate Fold 1-like conformations; however, both curves remained stable, suggesting that the trajectory remained close to Fold 1 without reaching Fold 2. These results demonstrate that replica exchange overcomes the limitation of single-temperature sampling by combining high-temperature exploration with low-temperature refinement, thereby facilitating transitions between Fold 1 and Fold 2 conformational states.

We also evaluated the sensitivity of MSFold to the number of replicas *M* and the number of sampling steps *N*. Detailed parameter-sensitivity results are provided in the Supplementary Information.

### 2.5 The discrete structure-token space reshapes conformational sampling

Having observed transitions between conformational states during MSFold sampling, we next examined how these transitions are represented in structure-token space. Previous applications of replica exchange for conformational exploration have primarily operated in continuous Cartesian coordinate space, whereas MSFold performs sampling in a discrete structure-token space. We therefore analyzed structure-token distributions and updates in LAOBP and examined how they are associated with conformational changes.

Fig. 4a shows the probability distributions of structure tokens predicted by ESM3 across LAOBP residues. Sampling from these distributions generates structure-token sequences that can be decoded into corresponding conformations. Some residues, such as Ile67 and Ile81, exhibit highly concentrated distributions dominated by only a few tokens. In contrast, residues 89–92 show broader distributions, substantially higher entropy, and a larger number of observed structure-token types. Notably, these residues form a strand that connects two lobes and constitutes a hinge whose backbone torsional rearrangements mediate the open-to-closed domain motion of LAOBP [28]. This uneven entropy profile indicates that token-level variability is not uniformly distributed across the sequence.

**Fig. 4:**
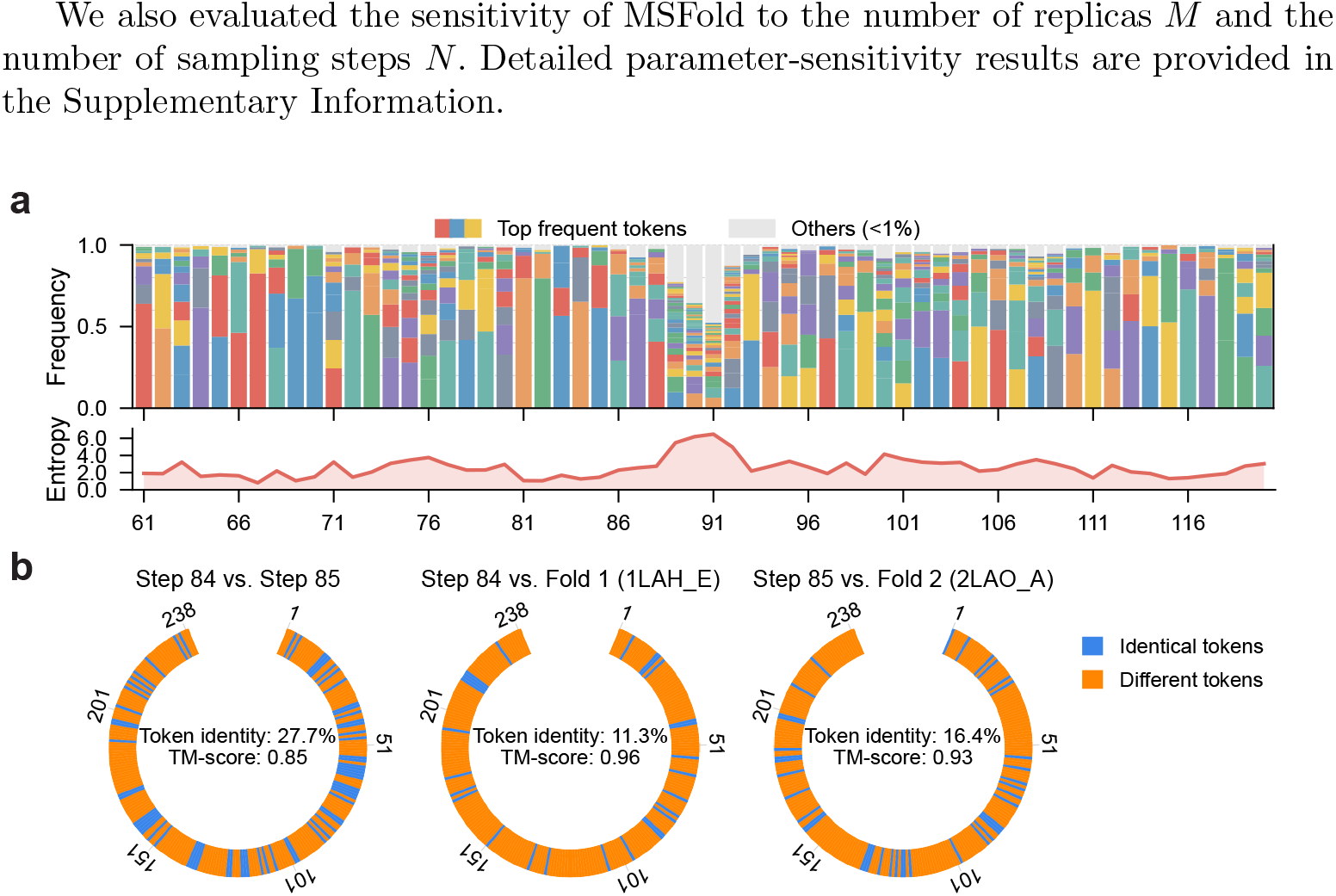
Distribution of structure tokens across LAOBP residues and large-scale token updates during MSFold sampling. **a,** Some residues, such as Ile67 and Ile81, exhibit highly concentrated distributions dominated by only a few tokens, whereas residues 89–92 show broader distributions with substantially higher entropy. **b,** At the transition from step 84 to step 85, MSFold updates 72.3% of the structure tokens, and the two conformations have a TM-score of 0.85. This transition illustrates how large-scale token updates within a single MSFold step can produce substantial conformational changes.

We next quantified the fraction of structure tokens updated per MSFold sampling step and measured the resulting conformational changes. As shown in Fig. 4b, individual sampling steps often updated more than 60% of residue-level structure tokens. A representative transition further showed that a rapid state switch can be accompanied by extensive reassignment of structure tokens. At the transition from step 84 to step 85, 72.3% of the structure tokens were updated. These results show that a single MSFold step can involve extensive structure-token reassignment accompanied by a substantial conformational change.

Comparisons with native references show a related pattern. Step 84 remains highly similar to the closed-state reference 1LAH (TM-score = 0.96) despite sharing only 11.3% token identity, whereas step 85 remains highly similar to the open-state reference 2LAO (TM-score = 0.93) despite sharing only 16.4% token identity. These observations show that discrete structure-token assignments can differ substantially even when the decoded structures remain close to native conformations. Conformations closely resembling a reference state can therefore correspond to substantially different structure-token sequences.

To complement the structure-token sampling results, we compared MSFold with atomistic replica-exchange molecular dynamics using REST2 (REST2-REMD) for three representative proteins (Supplementary Fig. S5). Under the simulation settings and timescale examined here, the REST2 trajectories predominantly remained close to their respective starting conformations, whereas MSFold generated conformations close to both experimentally determined reference states. Thus, in this focused comparison, MSFold showed broader coverage of the two reference conformations than the corresponding short REST2 simulations did.

Together, these results show that conformational states in MSFold are not associated with a single fixed structure-token sequence: native-like conformations can be retained despite substantial differences in token assignments, and transitions between conformational states can be accompanied by extensive token reassignment. The REST2 comparison further shows that, under the tested conditions, MSFold and short Cartesian-space simulations exhibit different sampling behavior and reference-state coverage.

### 2.6 Sequence log-likelihood improves selection of plausible conformations

After generating sampled candidate conformations, a critical step is selecting plausible conformations when no experimentally determined structures are available in advance. To aid the selection, we assigned each sampled candidate conformation a confidence score, termed sequence log-likelihood (SLL), which measures sequence–structure consistency (see Methods for details). We evaluated SLL’s ability to rank and select plausible conformations and compared it with pTM and pLDDT, the confidence scores used by AlphaFold3.

We first investigated the relationship between these confidence scores and structural accuracy using the sampled candidate conformations of the 1YNL L/2CGR L pair as a case study. As shown in Fig. 5a, SLL achieves a Spearman correlation coefficient of 0.71 with TM-score, higher than those of pTM (0.65) and pLDDT (0.68). Fig. 5b further shows two representative conformations. One resembles Fold 1 with a TM-score of 0.86 and ranks within the top 0.8% of the sampled candidate conformations according to TM-score, whereas the other resembles Fold 2 with a TM-score of 0.98 and ranks within the top 0.4%. Both conformations are ranked within the top 1% by SLL. In contrast, their rankings drop to the top 69% and 63%, respectively, according to pTM, and to the top 44% and 57% according to pLDDT. In this case, SLL more effectively prioritizes conformations resembling the experimentally determined structures than pTM and pLDDT.

**Fig. 5:**
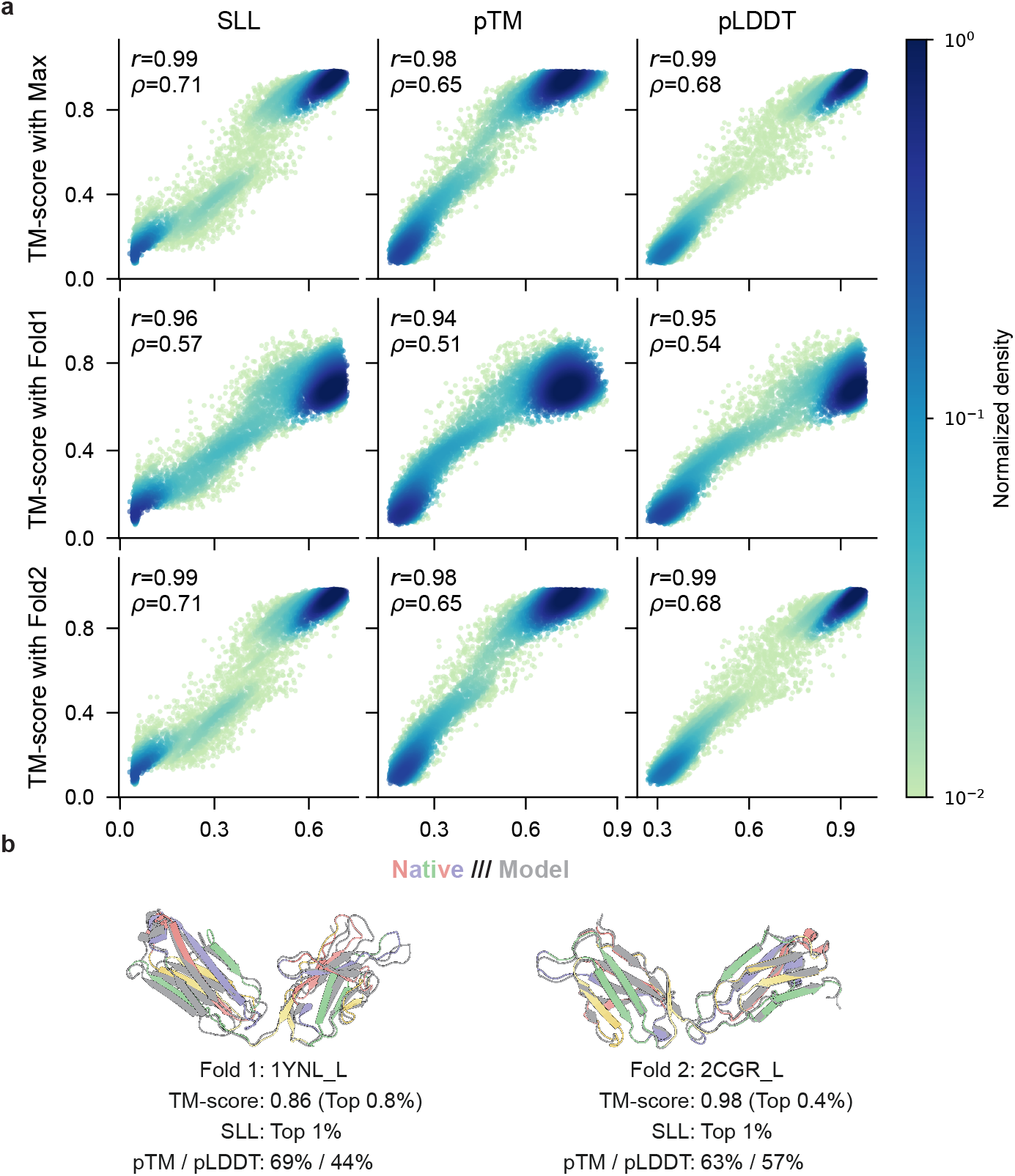
Evaluation of confidence scores for structural accuracy correlation and conformation ranking. **a,** Correlation between confidence scores and TM-score across the 20,000 conformations sampled for the 1YNL L/2CGR L pair. The figure reports the Pearson correlation coefficient *r* and Spearman correlation coefficient *ρ*. SLL shows a higher Spearman correlation with TM-score than pTM and pLDDT. **b,** Ranking of two representative conformations by the three confidence scores. The two conformations closely resemble Fold 1 and Fold 2, respectively. Both are ranked within the top 1% by SLL. In contrast, their rankings fall to the top 69% and 63%, respectively, according to pTM, and to the top 44% and 57%, respectively, according to pLDDT.

We further applied SLL to select plausible conformations across the 312 test protein pairs. As shown in Extended Data Table 2, the best-matching conformations in the full set of sampled candidate conformations achieve average TM-scores of 0.821 and 0.740 against Fold 1 and Fold 2, respectively. The top-ranked conformations selected by SLL achieve average TM-scores of 0.720 and 0.611, compared with 0.715 and 0.607 for pTM and 0.697 and 0.593 for pLDDT. We next investigated the best structural accuracy achieved among the top-*k* conformations ranked by each confidence score. Similar trends are observed for the top 5, 10, and 20 conformations, with SLL achieving slightly higher average TM-scores than pTM and pLDDT across these selections.

Overall, these results suggest that SLL provides a reference-free criterion for prioritizing sampled candidate conformations, with slightly better overall ranking performance and more pronounced advantages in some cases.

## 3 Discussion

Our results show that the multi-conformation prediction capability of a pretrained protein language model depends not only on what structural information the model has learned, but also on how that information is accessed at inference time. MSFold treats the structure-token space learned by ESM3 as a sequence-conditioned conformational search space and explores it through parallel tempering. This strategy improves recovery of Fold 2 conformations while preserving high accuracy for Fold 1 conformations, even for temporally held-out proteins released after the ESM3 training-data cutoff. These findings suggest that the pretrained model already encodes structural information relevant to multiple conformations, but standard decoding and local sampling strategies access only part of this information. Inference-time search therefore provides a complementary means of accessing the predictive capability of pretrained protein models without retraining.

A central implication of our results is that having a learned structure-token space alone is not sufficient for effective multi-conformation prediction; how this space is explored at inference time is also important. In this study, ESM3 provides a discrete structure-token space for conformational sampling, whereas MSFold uses parallel tempering to explore this space at inference time. Our trajectory analysis shows that a single sampling step can involve reassignment of a large fraction of structure tokens together with a transition between conformational states. Sampling in structure-token space also provides an alternative way to explore protein conformational space. The comparison with short REST2 simulations further illustrates differences in sampling behavior between MSFold and Cartesian-space simulations under the tested conditions.

At the same time, several limitations should be made explicit. The training-free nature of MSFold is both a strength and a constraint: it enables broader exploration of the learned structure-token space without retraining, but its performance remains bounded by the structural information encoded by the pretrained model. In addition, quantization and decoding errors in the discrete-structure representation impose an upper bound on the achievable structural fidelity. Finally, broader exploration incurs additional computational cost relative to standard structure-token sampling.

Broad exploration also raises another challenge: selecting plausible conformations from the sampled ensemble. Sequence log-likelihood provides a reference-free criterion for prioritizing conformations, with slightly better overall ranking performance and more pronounced advantages in some challenging cases. However, reliable selection of plausible alternative conformations remains unresolved. Future improvements will therefore likely require progress in both conformational exploration and selection.

Taken together, the results presented in this study establish MSFold as an effective approach for protein multi-conformation prediction. More broadly, they highlight a general principle for generative structure prediction: a pretrained protein language model may encode more structural information than standard decoding reveals, and realizing this latent capability depends on how the learned structural space is explored and how the resulting conformations are ranked.

## 4 Methods

The MSFold sampling procedure and sequence log-likelihood (SLL) confidence score are described below. Additional algorithmic details, together with benchmark dataset construction, baseline methods, evaluation metrics, and molecular dynamics simulations, are provided in the Supplementary Information.

### 4.1 Benchmark dataset and evaluation

We constructed a benchmark dataset comprising 312 protein pairs with experimentally determined multiple conformations. The benchmark was assembled from three sources: apo–holo protein pairs [23], fold-switching proteins [24], and a custom-curated subset of the Protein Data Bank (PDB) [6]. The custom PDB subset was constructed from structures available up to December 2022. Across the benchmark, proteins were restricted to 50–500 residues, sequence coverage was required to exceed 95%, paired states were required to have at least 95% sequence identity, and conformationally distinct pairs were retained with TM-score *<* 0.8; disordered proteins were excluded using DSSP-based filtering [29, 30].

We compared MSFold with AlphaFold 2 [10], AlphaFold 3 [11], MSA Subsampling [13], AF Cluster [14], ConfDiff [31], Str2Str [16], EigenFold [15], AlphaFlow [17], ESM3 with argmax decoding and iterative structure-token sampling [21], and ESMDiff [26].

Structural accuracy was evaluated using TM-score against each experimentally determined conformation [25]. Fold 1 denotes the conformation preferentially predicted by ESM3, whereas Fold 2 denotes the alternative conformation. This assignment is model-relative rather than biologically intrinsic. For each reference conformation, we report the maximum TM-score achieved by any generated candidate for that target. A target was considered successfully predicted only if both conformations were recovered with TM-score ≥ 0.75. Thus, the benchmark evaluates whether the generated ensemble contains accurate representatives of both experimentally observed conformational states.

We further conducted a temporal evaluation using the ESM3 training-data cutoff of 1 May 2020 [21]. The temporally held-out subset comprised 33 protein pairs whose experimentally determined conformations were released after this cutoff. We used the ESM3 cutoff rather than the latest cutoff across all comparison methods because applying the latter would leave no eligible evaluation cases. Because methods trained with later cutoffs may have had access to some of these structures, this analysis provides a conservative test of whether the advantage of MSFold can be explained by memorization of structures available during ESM3 training.

For a complementary analysis of whether prior availability of both reference conformations improves multi-conformation prediction, we used AlphaFold 2 as an example and selected 238 benchmark pairs for which both experimentally determined conformations were released before the AlphaFold 2 training-data cutoff of 30 April 2018 [10].

### 4.2 MSFold sampling in the discrete structure-token space for multi-conformation prediction

MSFold applies parallel tempering [22] to the sequence-conditioned structure-token space learned by ESM3 [21]. For each of the *M* replicas, ESM3 predicts position-wise structure-token distributions conditioned on the input sequence, from which an initial structure-token state is sampled independently. The replicas are assigned distinct temperatures along a temperature ladder, with higher-temperature replicas promoting broader exploration and lower-temperature replicas preserving more concentrated sampling of high-probability structural states.

Within each replica, MSFold performs blocked masked-token updates. During each sampling step, residue positions are visited in randomized order. For a target residue, a local block is defined using the target residue together with its spatially nearest neighbors in the current decoded backbone. The structure tokens within this block are masked simultaneously, and ESM3 predicts their conditional logits given the input sequence and the remaining token context. For replica *i* at temperature *T_i_*, the masked block *b* is resampled from the temperature-scaled distribution

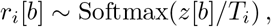

where *z*[*b*] denotes the ESM3 structure-token logits for the masked positions. Block updates allow coupled local structural changes to be proposed while retaining the sequence-conditioned structural prior learned by ESM3.

After each within-replica sweep, MSFold attempts local exchanges between adjacent-temperature replicas using an odd–even pairing schedule. Rather than exchanging complete replica states, an exchange region *I*_exchange_ is constructed from selected local blocks, and only the structure tokens within this region are proposed for exchange. Because ESM3 provides conditional structure-token distributions rather than an explicit normalized joint probability over complete token configurations, MSFold uses a Metropolis-style local exchange criterion based on conditional token pseudo-likelihoods. For adjacent replicas *i* and *j*, the current and proposed local configurations are assigned pseudo-likelihoods *P*_current_ and *P*_swap_, respectively, and the proposal is accepted with probability

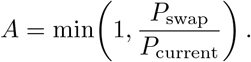

Here, *A* is the acceptance probability of the current exchange proposal; the realized accept/reject outcome is recorded separately. Local exchange allows configurations explored at higher temperatures to propagate toward lower-temperature replicas without requiring low-probability exchanges of complete high-dimensional token states. The temperature ladder is adapted during sampling because an appropriate spacing between adjacent temperatures is target-dependent. Following adaptive parallel-tempering schemes [32, 33], MSFold updates the log-gap variable Δ[*i*] for an exchanged replica pair according to the difference between the current exchange acceptance probability *α*_curr_[*i*] and a target probability *α*_target_,

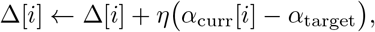

where the step size *η* decreases over the course of sampling. The updated positive gaps are then used to reconstruct the temperature ladder while keeping the minimum temperature fixed.

In this study, we used *M* = 40 replicas and *N* = 500 sampling steps, yielding *N × M* = 20,000 sampled structure-token states per target. The sampled token sequences were decoded into backbone coordinates using ESM3, and full-atom structures were reconstructed with FlowPacker [34]. The resulting conformations were ranked using the sequence log-likelihood described below.

### 4.3 Sequence log-likelihood for conformation ranking

To rank the conformations sampled by MSFold, we define a reference-free confidence score termed sequence log-likelihood (SLL), based on sequence–structure consistency. For a sampled conformation represented by its structure-token state **s**, ESM3 is used in inverse-folding mode to evaluate the probability assigned to the target amino acid at each residue position conditioned on the sampled structure-token state [21]. We define SLL as the average per-residue log-likelihood of the target sequence,

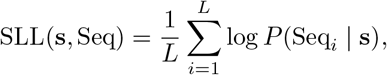

where *L* is the number of amino-acid residues in the target sequence.

Higher SLL values indicate greater consistency between the sampled structural representation and the target amino acid sequence. SLL is calculated after conformational sampling and is used only to rank the decoded conformations; it does not affect the structure-token sampling procedure or replica-exchange dynamics.

## Declarations

### Data availability

Source data are available on Zenodo at https://doi.org/10.5281/zenodo.20437832.

### Code availability

The MSFold code is available at https://github.com/findless/MSFold.

### Author contributions

Z.W. conceived and developed the method, performed the experiments and analyses, and drafted the manuscript. Y.Y. contributed to benchmark dataset construction. W.-M.Z., C.Y. and D.B. supervised the study and contributed to methodological discussions. D.B. also contributed to revision of the manuscript.

