## supplementary for "Sampling in structure-token space enables accurate prediction of multiple protein conformations"

### Supplementary Information for:
Sampling in structure-token space enables accurate prediction of multiple protein conformations

Zhiyuan Wang et al.

**2 Parameter sensitivity analysis** **3**

**3 Correlation analysis of token identity, embedding similarity, and structural similarity** **5**

**4 Two additional case studies of MSFold conformational sampling** **7**

**5 Cartesian-space conformational sampling using molecular dynamics** **9**

**Extended Data Tables** **11**

**Supplementary Methods** **12**

**6 Details of the MSFold algorithm** **12**

### Supplementary Results

#### 1 Construction of benchmark dataset, evaluated methods, and evaluation protocol

##### 1.1 Construction of benchmark dataset

We constructed a benchmark dataset comprising 312 protein pairs with experimentally determined multiple conformations. The benchmark was assembled from three sources: apo–holo protein pairs [1], fold-switching proteins [2], and a custom-curated subset of the Protein Data Bank (PDB) [3]. The custom PDB subset was constructed from structures available up to December 2022 and was designed to enrich for proteins with multiple experimentally resolved conformations while minimizing sequence differences between paired states.

For the custom PDB subset, structures were first filtered by sequence length (50–500 residues) and sequence coverage ($> 95\%$). Protein sequences were then clustered using MMseqs2 [4] at 100% sequence identity. Within each sequence cluster, US-align [5] was used to identify conformationally distinct structure pairs with TM-score $< 0.8$. Disordered proteins were further excluded using DSSP-based filtering [6, 7]. We applied the same basic criteria to the apo–holo and fold-switching subsets, excluding proteins outside the 50–500-residue range, pairs with conformational TM-scores $\geq 0.8$, and pairs with sequence identity below 95%.

After combining the three sources and applying these filters, the final benchmark contained 312 multi-conformation protein pairs. The distributions of protein length and conformational TM-score in the benchmark are shown in Fig. S1.

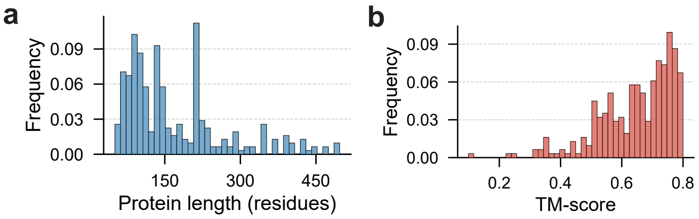

Figure S1: **Statistics of the multi-conformation benchmark dataset. a,** Distribution of protein sequence lengths in the benchmark dataset. **b,** Distribution of TM-scores between the two experimentally determined conformations (Fold 1 and Fold 2) for each protein pair.

##### 1.2 Evaluated methods for comparison

We compared MSFold with a diverse set of existing methods for a comprehensive comparison. We ran these methods with their default settings to yield predicted structures. Notice that the structures generated by these methods differ in count due to their distinct default settings, computational cost budgets, and method characteristics (e.g., MSA clustering). The methods and their specific configurations are listed as follows:

- AlphaFold 2: 25 structures were generated by its five internal models[8].
- AlphaFold 3: 25 structures were generated across five random seeds [9].
- MSA Subsampling: 10 structures were obtained from 10 independent runs with `max_msa_clusters = 32` [10].
- AF Cluster: 10 structures were obtained from 10 independent runs with `min_eps = 3` [11].
- ConfDiff: 200 structures were generated under its default setting [12].

- Str2Str: 200 structures were generated under its default setting [13].
- EigenFold: 5 structures were generated under its default setting [14].
- AlphaFlow: 10 structures were generated under its default setting [15].
- ESM3: 100 structures were produced with argmax selection and 100 with iterative structure-token sampling [16].
- ESMDiff: 100 structures were generated using 25 denoising steps [17].

##### 1.3 Evaluation protocol

Structural accuracy was evaluated using TM-score against each experimentally determined conformation [18]. Following the convention used throughout the main text, Fold 1 denotes the conformation preferentially predicted by ESM3, whereas Fold 2 denotes the alternative conformation. This assignment is therefore model-relative rather than biologically intrinsic, and performance is reported separately for the two states. A target was considered successfully predicted only if both conformations were recovered with TM-score $\geq 0.75$.

For each method, conformational coverage was evaluated at the ensemble level. For each reference conformation, we report the maximum TM-score achieved by any generated candidate for that target. The benchmark therefore measures whether the generated set of sampled candidate conformations contains accurate representatives of both experimentally observed conformational states. Reference-free identification of plausible conformations from sampled candidate conformations is evaluated separately in the ranking analysis described in the main text.

We further conducted a temporal evaluation to assess whether MSFold's performance extends beyond the structures available during ESM3 training. To this end, we constructed a temporally held-out subset comprising 33 protein pairs whose experimentally determined conformations were released after the ESM3 training-data cutoff of 1 May 2020. We used the ESM3 cutoff rather than the latest cutoff across all comparison methods because applying the latter would leave no eligible evaluation cases. The conformations in this subset were therefore unavailable in the structural data released before the ESM3 cutoff, although methods trained with later cutoffs may have had access to some of these structures. Consequently, this evaluation provides a conservative test of whether the advantage of MSFold can be explained by memorization of structures available during ESM3 training.

##### 1.4 Evaluation results on the benchmark dataset

Supplementary Table S1 reports the full per-method TM-score comparison corresponding to the benchmark summary in the main text.

#### 2 Parameter sensitivity analysis

We evaluated MSFold on the same 10 randomly sampled multi-conformation protein pairs used for the ablation experiments under a range of parameter settings to examine its sensitivity to the number of replicas $M$ and the number of sampling steps $N$. Across the tested numbers of replicas, Fold 1 and Fold 2 TM-scores remained relatively stable. Increasing the number of sampling steps generally led to modest improvements in both Fold 1 and Fold 2 TM-scores. These results indicate that MSFold remains effective across the tested parameter settings and does not depend on a narrowly tuned choice of the number of replicas or sampling steps.

Table S1: **Performance comparison on the multi-conformation benchmark.** The table reports the average TM-scores for Fold 1 and Fold 2, shown as Fold 1/Fold 2. The highest average TM-score for each fold is highlighted in bold.

| Method | Custom PDB subset | Apo-holo | Fold-switch | All |
|---|---|---|---|---|
| MSFold (Our method) | 0.820/**0.742** | **0.868**/**0.772** | 0.749/**0.652** | 0.821/**0.740** |
| AlphaFold 2 | 0.819/0.681 | 0.858/0.693 | 0.821/0.589 | 0.824/0.677 |
| MSA Subsampling | 0.787/0.670 | 0.832/0.686 | 0.766/0.643 | 0.790/0.670 |
| AF Cluster | 0.794/0.698 | 0.849/0.748 | 0.794/0.627 | 0.800/0.699 |
| AlphaFold 3 | **0.825**/0.706 | 0.866/0.705 | **0.833**/0.625 | **0.830**/0.701 |
| ESM3 (Argmax selection) | 0.764/0.682 | 0.825/0.709 | 0.677/0.588 | 0.765/0.679 |
| ESM3 (Iterative sampling) | 0.793/0.706 | 0.836/0.725 | 0.700/0.608 | 0.792/0.702 |
| ConfDiff | 0.672/0.606 | 0.744/0.637 | 0.646/0.549 | 0.679/0.606 |
| Str2Str | 0.693/0.613 | 0.764/0.655 | 0.679/0.550 | 0.700/0.613 |
| ESMDiff | 0.783/0.696 | 0.842/0.734 | 0.701/0.599 | 0.784/0.694 |
| EigenFold | 0.762/0.663 | 0.804/0.674 | 0.695/0.561 | 0.763/0.657 |
| AlphaFlow | 0.786/0.689 | 0.843/0.698 | 0.753/0.615 | 0.790/0.685 |

Table S2: **Parameter sensitivity analysis of MSFold.** Performance of MSFold with different numbers of replicas and sampling steps on the same 10 randomly sampled multi-conformation protein pairs used for the ablation experiments.

| **(a) Replicas** | | | **(b) Sampling steps** | | |
|---|---|---|---|---|---|
| #replicas | Fold 1 | Fold 2 | #steps | Fold 1 | Fold 2 |
| 20 | 0.831 | 0.782 | 100 | 0.825 | 0.778 |
| 30 | 0.839 | 0.794 | 200 | 0.834 | 0.781 |
| 40 | **0.840** | 0.792 | 300 | 0.836 | 0.787 |
| 50 | 0.837 | **0.798** | 400 | 0.836 | 0.788 |
| – | – | – | 500 | **0.840** | **0.792** |

### 3 Correlation analysis of token identity, embedding similarity, and structural similarity

The transition analysis in the main text shows that discrete structure-token assignments can change substantially while the decoded structures remain similar. To investigate the representation-level basis of this behavior, we compared similarity at three levels: discrete token identity, continuous token-embedding similarity, and structural similarity measured by TM-score. The central question is how similarity in discrete structure-token assignments relates to similarity in three-dimensional structure.

ESM3 [16] reconstructs protein structures from the continuous embeddings associated with structure-token indices rather than from the discrete indices themselves. Each sampled conformation can therefore be represented at three related levels: three-dimensional coordinates, continuous token embeddings, and discrete token indices. A simple distance-preserving correspondence between discrete token identity and structural similarity would imply that conformations with highly dissimilar token assignments should also be structurally dissimilar. However, the transition examples in the main text suggest that this correspondence is not one-to-one: substantially different token assignments can decode to structurally similar conformations.

To characterize this relationship systematically, we analyzed pairwise token identity, embedding similarity, and TM-score. For Fig. S2, the analysis was performed on 1,000 structures sampled from the full pool of 20,000 conformations included in the released source data. As shown in Fig. S2, token identity is concentrated within a relatively narrow range of approximately 0–0.4, whereas embedding similarity and TM-score span a broader range and exhibit similar bimodal distributions. Token identity remains correlated with TM-score ($r = 0.84$–$0.89$), but the corresponding RMSE is relatively large (0.54), indicating that token identity alone provides an imprecise measure of structural similarity. By contrast, embedding similarity shows stronger correlations with TM-score ($r = 0.89$–$0.96$) and a substantially smaller RMSE (0.11), suggesting that the continuous embedding representation is more closely aligned with structural similarity.

Together, these results indicate that the discrete structure-token representation is not related to three-dimensional structure through a simple one-to-one or distance-preserving mapping. Although the discrete tokens retain substantial structural information, structural similarity cannot be inferred reliably from token identity alone. The continuous embedding representation provides a closer correspondence to three-dimensional structural similarity.

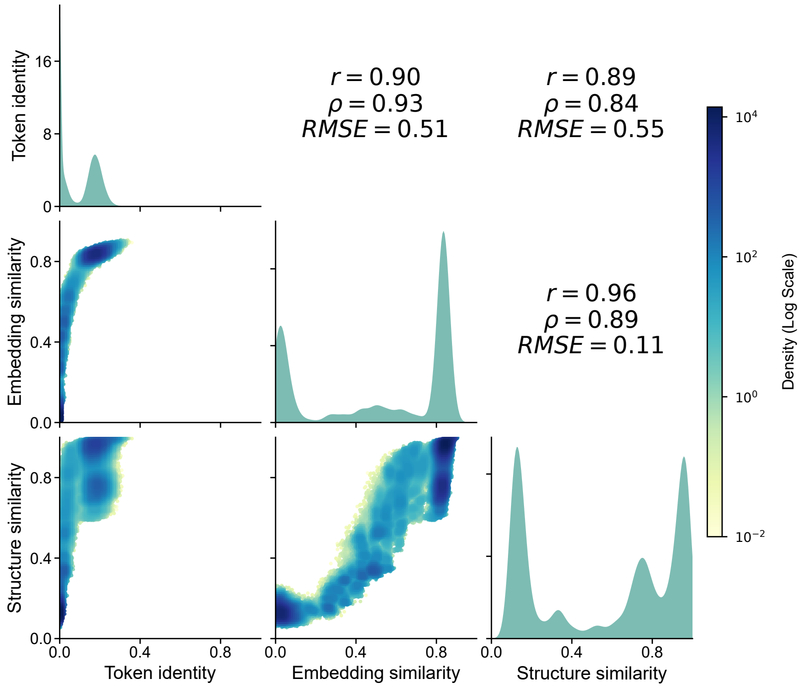

Figure S2: **Correlation analysis of token identity, embedding similarity, and structural similarity.** Pairwise token identity, embedding similarity, and structural similarity measured by TM-score were evaluated across a subset of 1,000 structures sampled from the full pool of 20,000 conformations. Diagonal panels show the marginal distributions of the three similarity measures, whereas off-diagonal panels show two-dimensional log-density histograms. Pearson correlation, Spearman correlation, and RMSE are reported in the mirrored panels. Token identity is concentrated within a relatively narrow range and shows a weaker correspondence with TM-score, whereas embedding similarity is more closely aligned with structural similarity.

### 4 Two additional case studies of MSFold conformational sampling

To assess whether the sampling behavior observed for the LAO-binding protein extends beyond a single target, we examined two additional benchmark systems: the 48G7 germline Fab and adenylate kinase. In both cases, MSFold accessed conformations close to both reference states within a single run and exhibited repeated transitions between the corresponding conformational regions.

For 48G7 (Fig. S3), the sampling trajectory alternates between conformations close to the two reference states, while the resulting ensemble forms two clusters centered around them. For adenylate kinase (Fig. S4), MSFold similarly accesses conformations close to both reference states and additionally samples intermediate conformations spanning the region between them. These additional examples indicate that the state-transition behavior observed for the LAO-binding protein is not restricted to a single benchmark case.

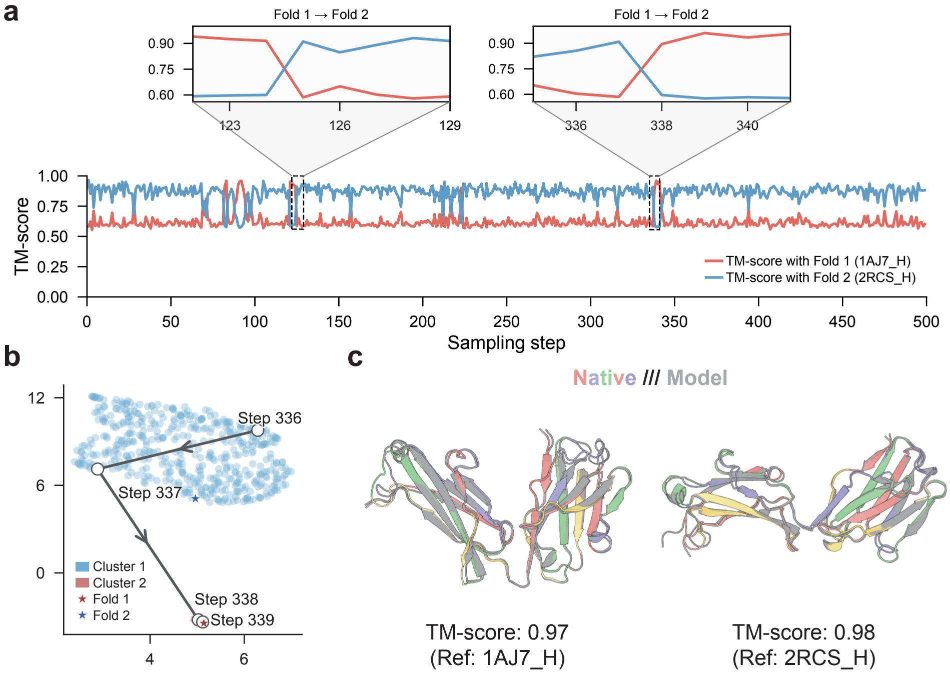

Figure S3: **MSFold sampling trajectory for protein 48G7. a,** Sampling trajectory measured by structural similarity (TM-score) to two reference conformations (Fold 1, PDB: 1AJ7_H, red; Fold 2, PDB: 2RCS_H, blue). In this example, the trajectory alternates between conformations close to the two reference states. Insets show representative transitions between the corresponding conformational regions during sampling. **b,** UMAP projection of the sampled conformational ensemble. Each point represents a sampled conformation, and two clusters are formed around the reference structures (stars). The arrowed interval marks the sampling segment shown in the right inset of **a** (steps 336–339). **c,** Structural superpositions of representative sampled models (grey) with the two reference structures (colored), with TM-scores of 0.97 and 0.98, respectively.

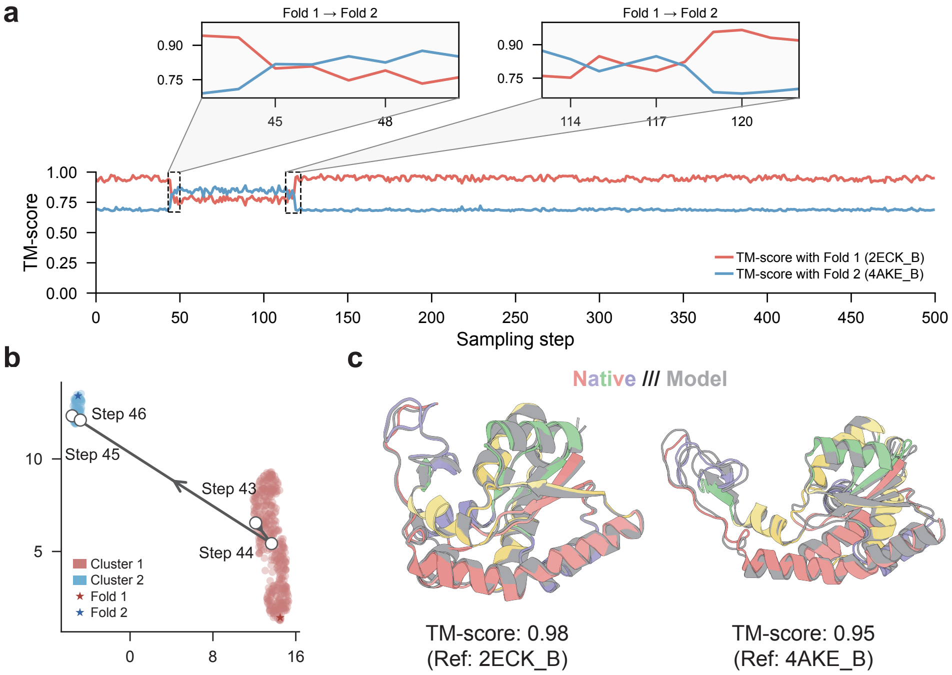

Figure S4: **MSFold sampling trajectory for adenylate kinase. a,** Sampling trajectory measured by structural similarity (TM-score) to two reference conformations (Fold 1, PDB: 2ECK_B, red; Fold 2, PDB: 4AKE_B, blue). In this example, the trajectory alternates between conformations close to the two reference states. Insets show representative transitions between the corresponding conformational regions during sampling. **b,** UMAP projection of the sampled conformational ensemble. Each point represents a sampled conformation, and two clusters are formed around the reference conformations (stars). The arrowed interval marks the sampling segment shown in **a** (steps 43–46). **c,** Structural superpositions of representative sampled models (grey) with the two reference structures (colored), with TM-scores of 0.98 and 0.95 for Fold 1 and Fold 2, respectively.

### 5 Cartesian-space conformational sampling using molecular dynamics

Previous applications of replica exchange for protein conformational exploration have primarily operated in continuous Cartesian coordinate space. Here, we used replica-exchange molecular dynamics as a Cartesian-space reference to examine the accessibility of alternative conformations for six trajectories across three representative proteins.

For each protein, replica-exchange molecular dynamics was performed using REST2 (REST2-REMD), with simulations initialized separately from Fold 1 and Fold 2. Protein-only systems were simulated using OpenMM/OpenMMTools with the Amber14 force field and TIP3P-FB solvent. Twenty replicas were distributed over an effective-temperature range of 300–450 K, and trajectories were analyzed at the demultiplexed 300 K state. Each replica was simulated for 10 ns. Because this timescale is short relative to many atomistic conformational transitions, these simulations were intended as a focused comparison of sampling accessibility rather than as a converged characterization of the underlying conformational dynamics. For each saved frame at 300 K, TM-scores to both experimentally determined reference conformations were calculated using the common set of protein residues.

Across the six REST2 trajectories, simulations predominantly remained close to their respective starting conformations (Supplementary Fig. S5). The trajectory initialized from 2RCS_H showed the strongest approach toward the alternative reference 1AJ7_H, reaching a maximum TM-score of 0.882, with 28.1% of analyzed frames achieving TM-score $\geq 0.75$. The remaining five trajectories did not exhibit a similarly sustained approach to the opposite reference state.

By contrast, MSFold sampling for the same three proteins generated conformations close to both experimentally determined reference states. Under the sampling conditions examined here, MSFold therefore achieved broader coverage of the two reference conformations than the corresponding short REST2 simulations. These results suggest that sampling in the learned structure-token space can improve access to alternative conformational states within the examined computational timescale.

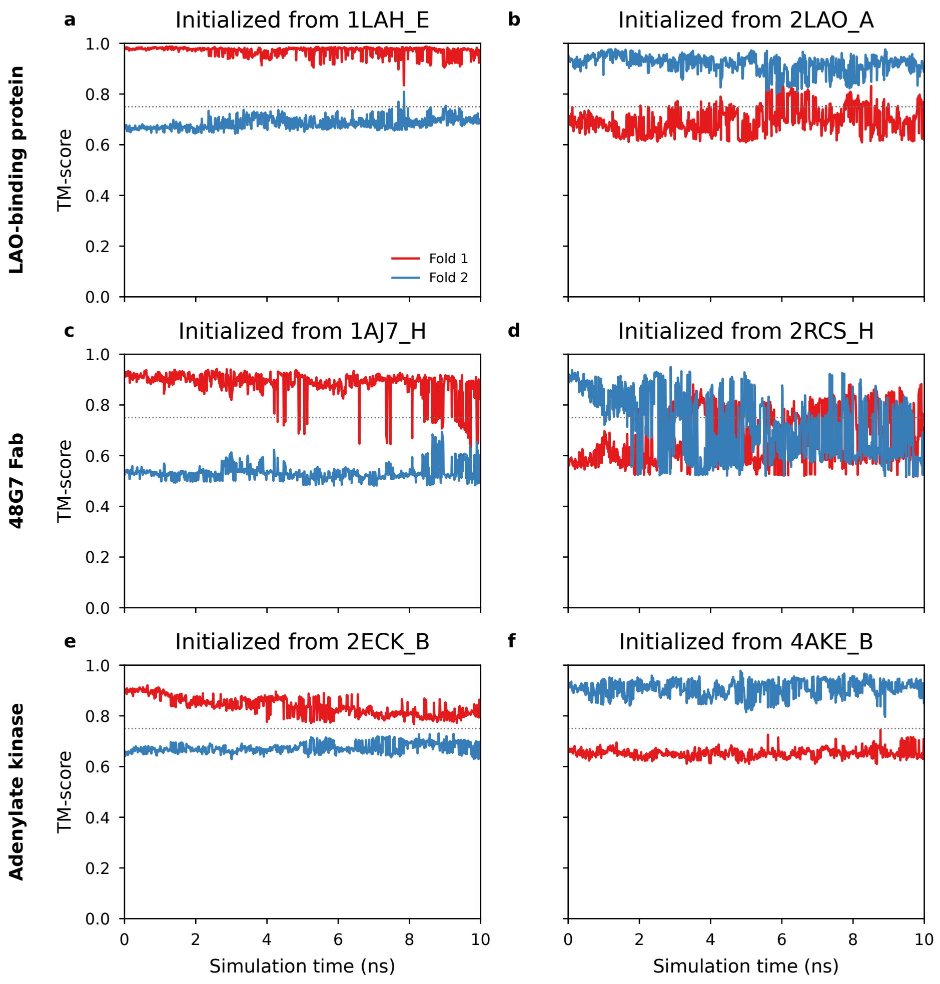

Figure S5: **Cartesian-space sampling with atomistic REST2 molecular dynamics.** TM-scores of demultiplexed 300 K REST2 frames relative to Fold 1 (red) and Fold 2 (blue). Each panel shows one 10-ns-per-replica trajectory initialized from the indicated experimental reference conformation. The horizontal dashed line marks TM-score = 0.75. The analysis includes three representative systems and is intended as a focused case-level comparison rather than a systematic benchmark of molecular dynamics.

### Extended Data Tables

**Extended Data Table 1 — Comparison of MSFold, its ablated variants, and ESM3 sampling strategies.** The methods were evaluated on 10 randomly sampled multi-conformation protein pairs. The two MSFold variants replace block updates with single-residue updates and disable replica exchange, respectively. The table reports the TM-scores of the best-matching sampled candidate conformations against Fold 1 and Fold 2 conformations.

| Method | Fold 1 | Fold 2 |
|---|---|---|
| **MSFold** | **0.840** | **0.792** |
| MSFold w/ single-residue updates | 0.823 | 0.792 |
| MSFold w/o replica exchange | 0.802 | 0.775 |
| ESM3 w/ argmax selection | 0.793 | 0.720 |
| ESM3 w/ iterative sampling | 0.794 | 0.746 |

**Extended Data Table 2 — Evaluation of confidence metrics for ranking sampled candidate conformations.** The table reports the TM-scores of the best-matching conformations in the full set of sampled candidate conformations against Fold 1 and Fold 2 conformations. It further reports the TM-scores of the best-matching conformations among the top-$k$ conformations ranked by each confidence metric. All TM-scores are averaged across the 312 protein pairs in the benchmark dataset.

| Metric | Best matching in sampled candidate conformations | TM-score of the best matching in top $k$ sampled candidate conformations (Fold 1/Fold 2) | | | |
|---|---|---|---|---|---|
| | | Top 1 | Top 5 | Top 10 | Top 20 |
| SLL | 0.821/0.740 | **0.720/0.611** | **0.749/0.639** | **0.757/0.648** | **0.766/0.657** |
| pTM | – | 0.715/0.607 | 0.743/0.635 | 0.755/0.644 | 0.762/0.652 |
| pLDDT | – | 0.697/0.593 | 0.731/0.627 | 0.743/0.639 | 0.752/0.648 |

### Supplementary Methods

#### 6 Details of the MSFold algorithm

This section provides pseudocode and details of the MSFold algorithm. The notation used throughout the algorithms is summarized in Table S3.

Table S3: Notations used by MSFold algorithm.

| Symbol | Description |
|---|---|
| **Core PT variables** | |
| $N_{\text{rep}}$ | Total number of replicas. |
| $N_{\text{iter}}$ | Total number of sampling iterations. |
| $t$ | Current sampling iteration index. |
| $\mathbf{T}$ | Vector of replica temperatures, with $T_1 > \cdots > T_{N_{\text{rep}}}$. |
| $T_i$ | Temperature of replica $i$. |
| $T_{\min}$ | Minimum (coldest) temperature. |
| $T_{\max}$ | Maximum (hottest) temperature. |
| $\boldsymbol{\Delta}$ | Vector of log-gap variables, where $\Delta[i]$ is the logarithm of the adjacent linear temperature gap between replicas $i$ and $i+1$. |
| $\alpha_{\text{target}}$ | Target exchange acceptance probability for adaptive temperature updates. |
| $\gamma_c$ | Baseline adaptation coefficient in the temperature-update rule. |
| $\gamma_\xi$ | Decay exponent for the adaptation step size. |
| $\Delta_{\max}$ | Upper bound on the log-gap variables. |
| $c_{\text{eff}}$ | Effective adaptation coefficient used at the current interval update. |
| **Protein and state variables** | |
| Seq | Target amino acid sequence. |
| $L$ | Sequence length, $L = \text{len(Seq)}$. |
| $\mathbf{R}$ | Matrix of current replica states of size $N_{\text{rep}} \times L$. |
| $r_i$ | Current structure-token state of replica $i$. |
| $\mathbf{s}$ | A sampled structure-token state. |
| $\mathbf{x}$ | Decoded three-dimensional structure for a sampled state $\mathbf{s}$. |
| $\mathcal{S}_{\text{raw}}$ | Set of collected raw structure-token states from all replicas. |
| $\mathcal{X}_{\text{set}}$ | Set of decoded structures. |
| $\mathcal{C}_{\text{set}}$ | Set of confidence scores. |
| $c$ | Confidence score for a sampled state. |
| **Sampling variables** | |
| $\mathbf{B}$ | Block map; $\mathbf{B}[j]$ is the local update block associated with residue $j$. |
| $N_{\text{nn}}$ | Number of nearest neighbors used to define a local block. |
| $b$ | A single local update block. |
| $N_{\text{exchange\_blocks}}$ | Number of sampled blocks used to define an exchange region. |
| $I_{\text{exchange}}$ | Set of token indices involved in an inter-replica exchange. |
| $\mathbf{z}$ | Structure-token logits predicted by ESM3 for masked positions. |
| $z_{\text{seq}}$ | Sequence logits predicted by ESM3 in inverse-folding mode. |
| $\boldsymbol{\alpha}_{\text{curr}}$ | Stored interval-specific acceptance probabilities used for temperature adaptation. |
| $A$ | Metropolis-style local exchange acceptance probability for a proposed exchange. |

Continued on next page

Table S3 – continued from previous page

| Symbol | Description |
|---|---|
| $\mathcal{P}$ | Set of adjacent replica pairs exchanged at the current iteration. |
| $\mathcal{I}$ | Set of temperature-gap indices updated at the current iteration. |
| **Likelihood variables** | |
| $L_{\text{total}}$ | Total log-likelihood of Seq given a sampled structure-token state $\mathbf{s}$. |

Table S4: **Default parameters used in MSFold.** Default values of the sampling and adaptive-temperature parameters used in the experiments.

| Paper symbol | Code parameter | Default value |
|---|---|---|
| $N_{\text{iter}}$ | `total_steps` | 500 |
| $N_{\text{rep}}$ | `temp_nums` | 40 |
| $T_{\text{min}}$ | `init_temp_min` | 0.5 |
| $T_{\text{max}}^{(0)}$ | `init_temp_max` | 10 |
| $c$ | `concentration` | 2.0 |
| $N_{\text{nn}}$ | `nn_nums` | 16 |
| $N_{\text{nn\_reset}}$ | `block_reset` | 25 |
| $N_{\text{exchange\_blocks}}$ | `swap_block_nums` | 3 |
| $\alpha_{\text{target}}$ | `alpha_star` | 0.234 |
| $\gamma_c$ | `gamma_c` | 0.25 |
| $\gamma_\xi$ | `gamma_ksai` | 0.6 |
| – | `max_interval` | 2.0 |
| $\Delta_{\text{max}}$ | derived from `max_interval` | $\log 2$ |

Here, $N_{\text{nn}} = 16$ denotes the target residue together with its 15 nearest neighbors. The parameter `max_interval`=2.0 is the upper bound on the linear temperature gap, whereas $\Delta_{\text{max}} = \log 2$ is the corresponding upper bound on the logarithmic gap. $T_{\text{max}}^{(0)}$ denotes the initial maximum temperature; adaptive updates may subsequently change the maximum temperature.

#### 6.1 Initialization of MSFold

Initialization in MSFold defines the current replica states, the temperature ladder, and the associated log-gap variables used for subsequent adaptive updates. For each of the $N_{\text{rep}}$ replicas, ESM3 predicts position-wise logits over structure tokens from the input sequence, and an initial token state is sampled independently from the resulting categorical distributions. This procedure yields a diverse set of initial states in structure-token space.

The second component of initialization is the temperature ladder $T_1 > T_2 > \cdots > T_{N_{\text{rep}}}$, with $T_1 = T_{\text{max}}^{(0)}$ and $T_{N_{\text{rep}}} = T_{\text{min}}$. The ladder must balance three requirements: $T_{\text{min}}$ should preserve sampling accuracy, $T_{\text{max}}^{(0)}$ should be high enough to facilitate transitions between local conformational regions, and adjacent temperatures must remain sufficiently close to permit effective exchanges. To allocate more sampling density to the low-temperature regime, where structural refinement is most important, we use a non-uniform schedule controlled by a concentration parameter $c = 2.0$. Specifically, we define

$$u_i = \left(\frac{i-1}{N_{\text{rep}}-1}\right)^c, \qquad i \in \{1, 2, \ldots, N_{\text{rep}}\}, \quad N_{\text{rep}} > 1,$$

and map these values to the logarithmic interval $[\log T_{\text{min}}, \log T_{\text{max}}^{(0)}]$ to obtain

$$T_i = \exp\Big(\log T_{\text{min}} + u_{N_{\text{rep}}-i+1}\big(\log T_{\text{max}}^{(0)} - \log T_{\text{min}}\big)\Big).$$

**Algorithm S1** MSFold algorithm

**Input:**

Seq: input amino acid sequence

**Output:**

$\mathcal{S}_{\text{raw}}$: collected structure-token states

$\mathcal{X}_{\text{set}}$: decoded structures

$\mathcal{C}_{\text{set}}$: confidence scores

▷ Step 1: Initialization

$(\mathbf{R}, \mathbf{T}, \boldsymbol{\Delta}) \leftarrow \text{InitializeReplicas}(\text{Seq})$ ▷ $\mathbf{R}$: current replica states

$L \leftarrow \text{len}(\text{Seq})$

$\mathbf{B} \leftarrow \text{PartitionBlocks}(\mathbf{R}, N_{\text{nn}})$

$\mathcal{S}_{\text{raw}} \leftarrow \emptyset$

▷ Step 2: Sampling loop

**for all** $t \in \{0, \ldots, N_{\text{iter}} - 1\}$ **do**

    **if** $t \pmod{N_{\text{nn\_reset}}} = 0$ **then**

        $\mathbf{B} \leftarrow \text{PartitionBlocks}(\mathbf{R}, N_{\text{nn}})$

    **end if**

    $\mathbf{R} \leftarrow \text{IntraReplicaUpdate}(\mathbf{R}, \text{Seq}, \mathbf{T}, \mathbf{B}, L)$ ▷ See Algorithm S2

    $(\mathbf{R}, \boldsymbol{\alpha}_{\text{curr}}) \leftarrow \text{InterReplicaExchange}(\mathbf{R}, \text{Seq}, \mathbf{T}, \mathbf{B}, t, L)$ ▷ See Algorithm S3

    **if** $t > 0$ **then**

        $(\mathbf{T}, \boldsymbol{\Delta}) \leftarrow \text{UpdateTemperatures}(\mathbf{T}, \boldsymbol{\Delta}, \boldsymbol{\alpha}_{\text{curr}}, t)$ ▷ See Algorithm S4

    **end if**

    $\mathcal{S}_{\text{raw}} \leftarrow \mathcal{S}_{\text{raw}} \cup \{\mathbf{R}[i] \mid i = 1..N_{\text{rep}}\}$

**end for**

▷ Step 3: Decode structures and evaluate confidence

$\mathcal{X}_{\text{set}}, \mathcal{C}_{\text{set}} \leftarrow \emptyset, \emptyset$

**for** each $\mathbf{s}$ in $\mathcal{S}_{\text{raw}}$ **do**

    $\mathbf{x} \leftarrow \text{DecodeTokensToCoords}(\mathbf{s})$

    $c \leftarrow \text{CalculateConfidence}(\mathbf{s}, \text{Seq})$ ▷ See Algorithm S5

    $\mathcal{X}_{\text{set}} \leftarrow \mathcal{X}_{\text{set}} \cup \{\mathbf{x}\}$

    $\mathcal{C}_{\text{set}} \leftarrow \mathcal{C}_{\text{set}} \cup \{c\}$

**end for**

**return** $\mathcal{S}_{\text{raw}}, \mathcal{X}_{\text{set}}, \mathcal{C}_{\text{set}}$

The corresponding interval variables $\Delta$ are initialized from this ladder as the logarithms of adjacent linear temperature gaps and are subsequently updated during adaptive temperature control. At initialization, $T_{\max}^{(0)}$ denotes the hottest replica temperature; after adaptation, the highest temperature is reconstructed from the updated gaps and may differ from $T_{\max}^{(0)}$, while $T_{\min}$ remains fixed. This parameterization keeps the initialization consistent with the later update rule, which adjusts $\Delta$ and reconstructs the ladder from $T_{\min}$.

#### 6.2 Intra-replica update of structural tokens

Within each replica, MSFold updates the current structure-token state by blocked masked-token sampling (Algorithm S2). A single sweep visits all $L$ residue positions in random order. For a target residue index $j$, the update block $b = B[j]$ contains residue $j$ together with its $N_{\mathrm{nn}} - 1$ spatially nearest neighbors in the current decoded backbone geometry, as defined by pairwise $C_\alpha$–$C_\alpha$ distances. This block design is motivated by the restricted local structural neighborhood used by the ESM3 encoder.

To update this block, the corresponding structure tokens are masked simultaneously while the remaining tokens are kept fixed. ESM3 then predicts logits for the masked positions conditioned on the input sequence and the unmasked token context. The new block state is sampled from the temperature-scaled categorical distribution

$$r_i[b] \leftarrow \mathrm{Sample}\big(\mathrm{Softmax}(z[b]/T_i)\big),$$

where $z[b]$ denotes the logits predicted for block $b$, $r_i$ denotes the current token state of replica $i$ and $T_i$ is its temperature.

We use block updates rather than single-residue updates because local conformational changes often arise from concerted variations across neighboring positions. Simultaneous resampling of a local neighborhood therefore provides more effective within-replica exploration by capturing these coupled local transitions, while remaining compatible with the conditional token prediction mechanism of ESM3.

#### 6.3 Inter-replica exchange of structural tokens

During inter-replica exchange, MSFold proposes local configuration swaps between adjacent-temperature replicas rather than exchanging their full global configurations (Algorithm S3). This localized design addresses the main difficulty of exchange in high-dimensional systems: proposals that swap full global configurations typically introduce overly large discrepancies between replicas and therefore lead to very low acceptance probabilities. Restricting the proposal to a local subset of positions preserves more configurational overlap and yields practical exchange probabilities.

At each iteration, exchange attempts are made between adjacent replica pairs using an odd–even schedule, as defined in Algorithm S3. For each selected pair, the exchange region $I_{\mathrm{exchange}}$ is constructed by first sampling $N_{\mathrm{exchange\_blocks}}$ residue indices and then taking the union of their corresponding local blocks. Only the structure tokens within this region are proposed for exchange, while the remainder of each replica state is held fixed.

The exchange acceptance probability is computed using a Metropolis-style local exchange criterion based on conditional token pseudo-likelihoods over this local region. Because ESM3 provides conditional structure-token distributions rather than an explicit normalized joint distribution over complete token configurations, this procedure should not be interpreted as exact sampling from a known Boltzmann distribution. For a selected adjacent pair $i$ and $j$, let $p_i$ and $p_j$ denote the conditional token distributions over $I_{\mathrm{exchange}}$ under temperatures $T_i$ and $T_j$, respectively. The current and swapped local configurations are then scored as

$$P_{\mathrm{current}} = P(r_i[I_{\mathrm{exchange}}] \mid p_i)\ P(r_j[I_{\mathrm{exchange}}] \mid p_j)\,,$$

$$P_{\text{swap}} = P(r_j[I_{\text{exchange}}] \mid p_i)\ P(r_i[I_{\text{exchange}}] \mid p_j)\,,$$

and the proposal is accepted with probability

$$A = \min\left(1, \frac{P_{\text{swap}}}{P_{\text{current}}}\right).$$

In practice, this comparison is evaluated in the log domain, consistent with Algorithm S3. Here, $\boldsymbol{\alpha}_{\text{curr}}[i]$ denotes the current per-proposal exchange acceptance probability used for temperature adaptation, whereas the realized exchange outcome is recorded separately as a binary indicator. The interval index $i$ in the paper is one-based and corresponds to implementation array index $i-1$. This local exchange scheme allows conformations discovered at higher temperatures to be transferred to lower-temperature replicas, thereby helping the system escape kinetic traps without requiring low-probability global exchanges.

#### 6.4 Adaptive temperature update

Efficient inter-replica exchange depends critically on the temperature ladder. Because the optimal spacing between adjacent temperatures is target-dependent and not known in advance, a fixed ladder may yield inefficient exchanges for a given target. To address this limitation, MSFold adopts an adaptive temperature-tuning scheme [19] that updates the log-gap variables $\Delta$ rather than keeping the temperature ladder fixed. Here, $\Delta[i]$ denotes the logarithm of the linear temperature gap between adjacent replicas, so that the ladder can be reconstructed while preserving positivity of the gaps. The objective is to maintain exchange acceptance probabilities near a target value while preserving sufficient temperature separation for broad exploration.

For each interval selected at the current iteration, the update is based on the discrepancy between the current exchange acceptance probability $\alpha_{\text{curr}}[i]$ and the target probability $\alpha_{\text{target}}$ [19, 20]:

$$\Delta[i] \leftarrow \Delta[i] + \eta \cdot \big(\alpha_{\text{curr}}[i] - \alpha_{\text{target}}\big),$$

where the step size decays with iteration number as

$$\eta = \frac{c_{\text{eff}}}{(t+1)^{\gamma_\xi}}.$$

To accelerate correction when exchange probabilities fall below the target, we use an asymmetric coefficient $c_{\text{eff}}$, defined as $(1-\alpha_{\text{target}})/\alpha_{\text{target}}$ when $\alpha_{\text{curr}}[i] < \alpha_{\text{target}}$ and as $\gamma_c$ otherwise. Thus, intervals with insufficient exchange are contracted more aggressively, whereas intervals with exchange probabilities above the target are adjusted more gradually.

A further difficulty is high-temperature drift. If the target acceptance probability is enforced too rigidly across the entire ladder, especially in the extreme high-temperature regime, the upper intervals may continue to widen and drive $T_{\text{max}}$ toward excessively high values. To stabilize the ladder, we therefore impose an upper bound $\Delta_{\text{max}}$ on the log-gap variables and use the asymmetric coefficient above to contract excessively large gaps more aggressively when acceptance falls below the target. After the selected intervals are updated and clamped, the temperature ladder is reconstructed from $T_{\text{min}}$ and the resulting positive adjacent gaps.

#### 6.5 Decoding sampled structure-token sequences and ranking conformations

After the sampling loop is completed, the structure-token states collected across replicas and sampling steps are aggregated into the raw sample set $S_{\text{raw}}$. Each sampled structure-token sequence is then decoded into backbone coordinates using ESM3 [16]. Although the original ESM3 study reported all-atom structure decoding, the publicly available `esm3-sm-open-v1` model provides backbone coordinates only. We therefore use FlowPacker [21] to reconstruct full-atom structures from the decoded backbones.

Confidence scores are computed for all sampled structure-token sequences after sampling, following the SLL definition in the Methods section of the main text and summarized in Algorithm S5. Specifically, each sampled token sequence in $S_{\text{raw}}$ is used both to decode the corresponding structure and to compute its confidence score. The resulting conformations are then ranked according to these confidence scores.

**Algorithm S2** Intra-replica update of structural tokens

**Input:**

$\mathbf{R}$: replica states

Seq: input amino acid sequence

$\mathbf{T}$: temperatures

$\mathbf{B}$: block map

$L$: sequence length

**Output:**

$\mathbf{R}$: updated replica states

**for all** replicas $i \in \{1, \ldots, N_{\text{rep}}\}$ **in parallel do**
    $\mathcal{O} \leftarrow \textsc{RandomPermutation}(\{1, \ldots, L\})$
    **for** each residue index $j$ in $\mathcal{O}$ **do**
        $b \leftarrow \mathbf{B}[j]$
        $\tilde{r}_i \leftarrow r_i$
        $\tilde{r}_i[b] \leftarrow \text{MASK}$
        $z \leftarrow \textsc{StructureTokenLogits}(\text{Seq}, \tilde{r}_i, b)$
        $r_i[b] \leftarrow \textsc{Sample}(\text{Softmax}(z[b]/T_i))$
    **end for**
**end for**
**return R**

**Algorithm S3** Inter-replica exchange of structural tokens

**Input:**

$\mathbf{R}$: replica states
Seq: input amino acid sequence
$\mathbf{T}$: temperatures
$\mathbf{B}$: block map
$t$: iteration index
$L$: sequence length

**Output:**

$\mathbf{R}$: updated replica states
$\boldsymbol{\alpha}_{\text{curr}}$: interval-specific exchange acceptance probabilities

$I_{\text{sel}} \leftarrow \textsc{SampleIndices}(\{1, \ldots, L\}, N_{\text{exchange\_blocks}})$
$I_{\text{exchange}} \leftarrow \bigcup_{j \in I_{\text{sel}}} \mathbf{B}[j]$
**if** $t$ is even **then**
  $\mathcal{P} \leftarrow \{(1,2),(3,4),(5,6),\ldots\}$
**else**
  $\mathcal{P} \leftarrow \{(2,3),(4,5),(6,7),\ldots\}$
**end if**
$\boldsymbol{\alpha}_{\text{curr}} \leftarrow 0$
**for all** $(i,j) \in \mathcal{P}$ **in parallel do**
  $\tilde{r}_i \leftarrow r_i$; $\tilde{r}_j \leftarrow r_j$
  $\tilde{r}_i[I_{\text{exchange}}] \leftarrow \text{MASK}$
  $\tilde{r}_j[I_{\text{exchange}}] \leftarrow \text{MASK}$
  $z_i \leftarrow \textsc{StructureTokenLogits}(\text{Seq}, \tilde{r}_i, I_{\text{exchange}})$
  $z_j \leftarrow \textsc{StructureTokenLogits}(\text{Seq}, \tilde{r}_j, I_{\text{exchange}})$
  $p_i \leftarrow \text{Softmax}(z_i/T_i)$
  $p_j \leftarrow \text{Softmax}(z_j/T_j)$
  $\log P_{\text{current}} \leftarrow \text{LogLikelihood}(r_i[I_{\text{exchange}}], p_i) + \text{LogLikelihood}(r_j[I_{\text{exchange}}], p_j)$
  $\log P_{\text{swap}} \leftarrow \text{LogLikelihood}(r_j[I_{\text{exchange}}], p_i) + \text{LogLikelihood}(r_i[I_{\text{exchange}}], p_j)$
  $A \leftarrow \min(1, \exp(\log P_{\text{swap}} - \log P_{\text{current}}))$
  $\boldsymbol{\alpha}_{\text{curr}}[i] \leftarrow A$
  **if** $\textsc{Uniform}(0,1) < A$ **then**
    $\textsc{exchange}(r_i[I_{\text{exchange}}], r_j[I_{\text{exchange}}])$
  **end if**
**end for**
**return** $\mathbf{R}, \boldsymbol{\alpha}_{\text{curr}}$

**Algorithm S4** Adaptive temperature update

**Input:**
$\mathbf{T}$: temperatures
$\boldsymbol{\Delta}$: log-gap variables
$\boldsymbol{\alpha}_{\text{curr}}$: current per-proposal exchange acceptance probabilities
$t$: iteration index
**Output:**
$\mathbf{T}$: updated temperatures
$\boldsymbol{\Delta}$: updated gap variables

**if** $t$ is even **then**
    $\mathcal{I} \leftarrow \{1, 3, 5, \dots\}$
**else**
    $\mathcal{I} \leftarrow \{2, 4, 6, \dots\}$
**end if**
**for** each $i$ in $\mathcal{I}$ **do**
    $\text{diff} \leftarrow \boldsymbol{\alpha}_{\text{curr}}[i] - \alpha_{\text{target}}$
    **if** $\text{diff} < 0$ **then**
        $c_{\text{eff}} \leftarrow (1 - \alpha_{\text{target}})/\alpha_{\text{target}}$
    **else**
        $c_{\text{eff}} \leftarrow \gamma_c$
    **end if**
    $\eta \leftarrow c_{\text{eff}}/(t+1)^{\gamma_\xi}$
    $\boldsymbol{\Delta}[i] \leftarrow \boldsymbol{\Delta}[i] + \eta \cdot \text{diff}$
    $\boldsymbol{\Delta}[i] \leftarrow \min(\boldsymbol{\Delta}[i], \Delta_{\max})$
**end for**
$\mathbf{T}[N_{\text{rep}}] \leftarrow T_{\min}$
**for** $i \leftarrow N_{\text{rep}} - 1$ down to 1 **do**
    $\mathbf{T}[i] \leftarrow \mathbf{T}[i+1] + \exp(\boldsymbol{\Delta}[i])$
**end for**
**return** $\mathbf{T}, \boldsymbol{\Delta}$

**Algorithm S5** Confidence calculation via average sequence log-likelihood

**Input:**
$\mathbf{s}$: sampled structure-token state
Seq: target amino acid sequence
**Output:**
$c$: confidence score

$z_{\text{seq}} \leftarrow \textsc{SequenceLogits}(\mathbf{s})$
$L_{\text{total}} \leftarrow \textsc{ComputeLogLikelihood}(z_{\text{seq}}, \text{Seq})$
$c \leftarrow L_{\text{total}}/\textsc{len}(\text{Seq})$
**return** $c$
